# *In-vivo* distributed multicellular control of gene expression in microbial consortia

**DOI:** 10.1101/2023.11.13.566807

**Authors:** Davide Salzano, Barbara Shannon, Claire Grierson, Lucia Marucci, Nigel J Savery, Mario di Bernardo

## Abstract

This paper presents an innovative approach in synthetic biology, focusing on the engineering of biomolecular feedback control systems within microbial consortia to achieve robust and precise regulation of desired phenotypes. Traditional biomolecular control strategies, while effective, are predominantly confined to single-cell applications, limiting their complexity and adaptability due to metabolic constraints. We propose a novel methodology that distributes control functionalities across different cell populations within a microbial consortium. This approach leverages the division of labor and cooperative interactions among microbial populations, significantly enhancing flexibility and robustness. We build on the foundation of synthetically engineered microbial consortia, known for their applications in complex compound production and advanced computational processes. However, the challenge of implementing a distributed feedback control loop for precise phenotype regulation in synthetic communities has remained unaddressed. Our work closes this gap by developing and validating an in-vivo feedback control loop distributed across a bacterial consortium, specifically designed using two Escherichia coli strains. These strains are engineered to perform the three essential control functions: computation, sensing, and actuation. We employ orthogonal quorum sensing signaling molecules for communication between the controller and target cell populations. Through in-vivo experiments, we compare closed-loop and open-loop configurations to demonstrate the effectiveness of our approach. The results show that our distributed control system can regulate gene expression levels in target populations with high precision and robustness, even in the presence of consortium composition variations. This feature is particularly vital for practical applications in synthetic biology where stability and adaptability are crucial. Our study not only addresses a significant gap in synthetic biology literature but also opens avenues for more complex and reliable synthetic biology applications.

## Introduction

Synthetic biology aims to engineer biomolecular systems to achieve new functions [1]; potential applications range from designing bacteria that can produce biofuels or sense and degrade pollutants in the environment (such as hydrocarbons and plastics), to immune cells that can track and kill cancer cells, or that can release drugs at specific points in desired conditions to avoid side effects (see [2] for an extensive review). Biomolecular feedback controllers have been highlighted as a crucial ingredient for reliable and robust synthetic biology devices, increasing their flexibility and modularity [3]–[5].

Different designs have been proposed to implement biomolecular feedback control strategies both in-silico and in-vivo, ranging from simple feedback and feedforward loops to PID (Proportional Integral Derivative) controllers (see [3] and references therein). In particular, the introduction of an integral control action via an antithetic feedback motif has been highlighted as a fundamental ingredient for a synthetic circuit able to achieve and maintain a desired output even in the presence of disturbances, a property that has been termed *robust perfect adaptation* [6]. Integral biomolecular controllers have been thoroughly studied both theoretically [7]–[9] and experimentally, and different implementations have been proposed in bacteria [10] and mammalian cells [11]. For a review of the recent advancements in the implementations of biomolecular controllers guaranteeing robust perfect adaptation see [4].

However, existing robust controllers predominantly function at the single-cell level, posing inherent limitations on circuit complexity due to metabolic burden, and hindering their application as flexible plug-and-play modules across diverse synthetic and endogenous systems [5]. To overcome these constraints, following [12], we propose to distribute the required functionalities across distinct cell populations within a microbial consortium. Each cell strain encapsulates specific engineered gene networks, forming the foundation for a distributed feedback control loop [13], [14] involving two populations: a controller population and a target cell population. The former senses the targets’ output, compares it with a reference signal encoding the desired expression level, and provides the latter with stimuli based on antithetic control logic so as to steer the sensed output towards the desired goal [12], [15], [16].

Our work builds upon the foundation of synthetically engineered microbial consortia, where microbial cooperation, segregation, and division of labor have been leveraged for applications ranging from complex compound production [17] to advanced computational processes [18] (refer to [19]–[21] for a comprehensive review). Despite these advancements, as highlighted in [4], the in vivo implementation of synthetic communities implementing a distributed feedback control loop for the robust regulation of a desired phenotype to a specific reference value remains unexplored. Prior studies, such as Shou’s [22], have established mutualistic interactions between yeast populations for coexistence, and research documented in [23] achieved synchronized oscillations in E. coli strains via an engineered feedback loop. Furthermore, a majority sensing mechanism in a microbial consortium was realized through negative feedback in [24]. While these studies successfully created synthetic microbial consortia with targeted phenotypes, they lack the capability for precise and robust tuning of expression, a critical aspect for advanced synthetic biology applications.

In this paper, we address this fundamental gap in the literature by developing and validating in-vivo a feedback control loop distributed across a bacterial consortium. We demonstrate its effectiveness in regulating a phenotype of interest in the target population even in the presence of variations in the consortium composition. This is achieved by designing two Escherichia coli strains embedding all the modules required to implement the three fundamental functions needed for control, namely computation, sensing and actuation [12]. We show that the designed cellular populations respond to their respective inputs and that the two populations can exchange information through two rationally designed communication channels, implemented using orthogonal quorum sensing signalling molecules. To demonstrate the validity of the proposed strategy, we carry out in-vivo experiments comparing the closed loop configuration, where the controller cells can sense the signalling molecules from the targets, with an open-loop implementation, where they cannot. Our results show that the proposed multicellular control approach can successfully be used to achieve reliable and robust regulation of the expression level of a gene of interest in the targets while guaranteeing robustness to composition imbalances in the consortium, a particularly important property for practical implementations.

## Results

### A multicellular distributed control architecture

The implementation of a multicellular control architecture requires the rational design of two populations, namely the *controllers* and the *targets* (see Figure 1a). The controllers receive information about the targets’ output (*y*) which is the expression level of some gene of interest therein. This value is compared to that of some reference signal (*ref*) encoding the desired expression level and a control input (*u*) is then generated via the designed control logic. This signal is transmitted to the targets in order to regulate the level of their output to that encoded by the reference. The communication between controllers and targets is implemented via orthogonal Quorum Sensing molecules, where each cell population produces signalling molecules that can diffuse through the growth environment and into the other population.

**Figure 1.**
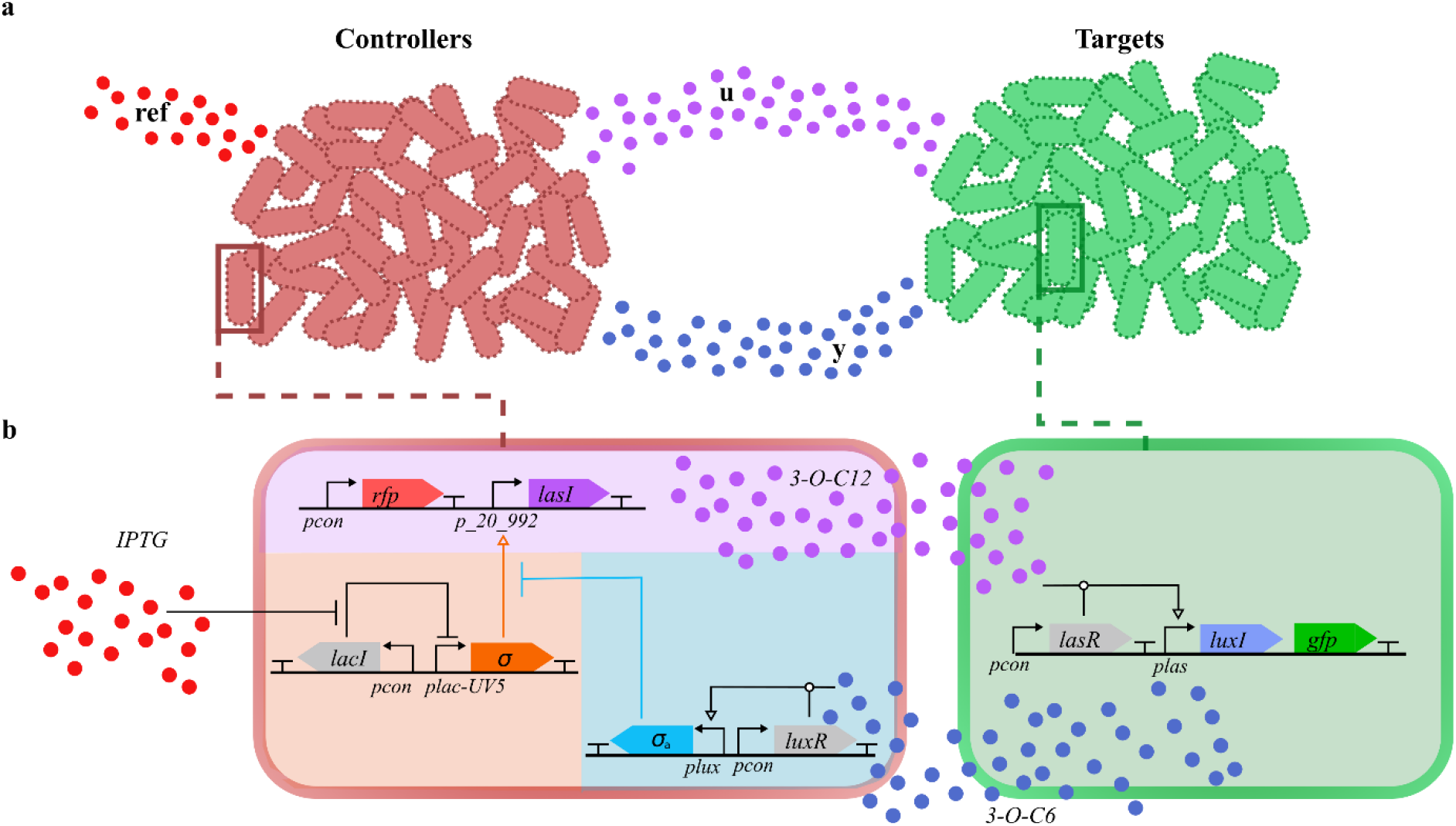
Multicellular control schematic. **A** Microbial consortium implementing a multicellular control architecture. The two populations, i.e. controllers and targets, share information using orthogonal quorum sensing channels (**u** and **y**), playing the role of the control input and the output of the process, respectively. In addition, the controllers sense the external stimulus labeled as **ref**, used to set up the reference of the feedback loop. **b**. Biological implementation of the multicellular control architecture. The Synthetic Biology Open Language (SBOL) notation is used to denote promoters, genes, promotion/inhibition relationships and quorum sensing molecules. The shaded areas identify different functional modules within each population.

The control logic, embedded in the controllers, is an antithetic feedback control strategy [3], based on the *in-silico* analysis presented in [12] and derived from the error computation module (comparator) implemented *in-vivo* using molecular titration in [25] and characterized dynamically in [26]. The controller network is a multi-input, single-output device that uses two independent signals to regulate the expression level of its output protein (see Figure 1b). It is constructed around a *σ* factor [27], which is a protein that recruits RNA polymerase (RNAP) to target promotors by binding to DNA sequences in the −10 and −35 regions of those promoters, and their cognate anti-*σ*, which blocks the interaction between RNAP and the *σ* factor by sequestering the *σ* factor. The production of *σ* is controlled by IPTG (the reference signal), which induces expression from the promoter *plac-UV5*. The production of anti-*σ* is regulated by the *plux* promoter, which is activated by the complex formed by LuxR, produced constitutively, and 3-O-C6-HSL, produced by the targets. The output signal of the controllers (3-O-C12-HSL) is produced by the product of the *lasI* gene, which is regulated by the *p20 992* promoter. This promoter, placed upstream of *lasI*, is induced by the presence of the orthogonal *σ* factor. The *σ* and anti-*σ* factors interact by binding and forming a complex that is unable to recruit RNAP to *p20_992*. All proteins are fused to a degradation tag (ssrA tag) [28].

The targets can express a green fluorescent protein (GFP), whose expression level we aim to regulate. The *gfp* gene is transcribed together with *luxI*, whose product is responsible for the production of the molecule 3-O-C-6-HSL. This small molecule operates as a proxy of the *gfp* expression levels by diffusing in the growth environment and activating the production of anti-*σ* in the controllers. The production of LuxI and GFP is regulated by *plas*. This promoter is activated by the complex formed by LasR, which is constitutively expressed, and 3-O-C12-HSL, which is produced by the controller population. In our implementation, LuxI was fused to a degradation tag (ssrA tag) to ensure its fast degradation. All the details of the plasmids and strains used in this work can be found in the *Strains and constructs* section of the Methods.

The goal of the design is to make the two engineered populations cooperate to steer and maintain *gfp* expression in the targets at a desired setpoint level, which can be changed by modifying the concentration of IPTG. Specifically, when *gfp* is overexpressed, the excessive production of 3-O-C6-HSL upregulates the production of anti-*σ*. In turn, a reduction of the free available *σ*, downregulates the production of LasI and, in turn, of the signalling molecule 3-O-C12-HSL. As a consequence, the activity of *pLas* in the targets should be reduced, decreasing GFP expression levels. Similar mechanisms lead to an increase in GFP when GFP falls below the desired level.

### Gene expression can be regulated via a multicellular architecture

Prior to mixing the two populations in the consortium, we tested the functional modules embedded in each population using flow cytometry [29]. Details on the machines and the settings used can be found in section *Samples preparation and analysis using flow cytometry* of the Methods.

We started by characterizing the response of the controller population to IPTG and 3-O-C6-HSL, which are used in the architecture as a reference signal and a proxy for the targets state, respectively. Note that, to measure the output level of the controllers, we substituted *lasI* with a *gfp* fused with an ssrA tag (see Figure S1). We grew the controllers at 37 °C for 3h adding different concentrations of IPTG and 3-O-C6-HSL to the growth media to test their response to their inputs. The fluorescence levels of the population was then analysed by Flow Cytometry (for more details see section *Induction protocol* of the Methods). The addition of IPTG promoted *plac-UV5* activity by inhibiting the repression exerted by LacI, thus inducing up to 300 fold increase in the steady state fluorescence when 100 *μM* IPTG was added to the media (Figure 2a). Conversely, the addition of 3-O-C6-HSL increased the anti-*σ* level of transcription by activating *plux;* as a consequence, the fluorescence levels consistently decreased, resulting in the inhibition of *gfp* expression levels up to 100 folds when 100 nM 3-O-C6-HSL was added to the growth media (Figure 2a). We used the same procedure to assess the response of the targets to 3-O-C12-HSL; as expected, addition of 10 *μM* 3-O-C12 in the growth media promoted *plas* activity and a 14 fold increase was detected in average GFP fluorescence (Figure 2b). These assays confirmed that the controllers were correctly responding to changes either in the reference signal or in the output of the targets, and, confirmed that the targets were able to adjust GFP production according to the concentration of 3-O-C12-HSL. For further details on the standard deviation of the response of targets and controllers see Figure S2.

**Figure 2.**
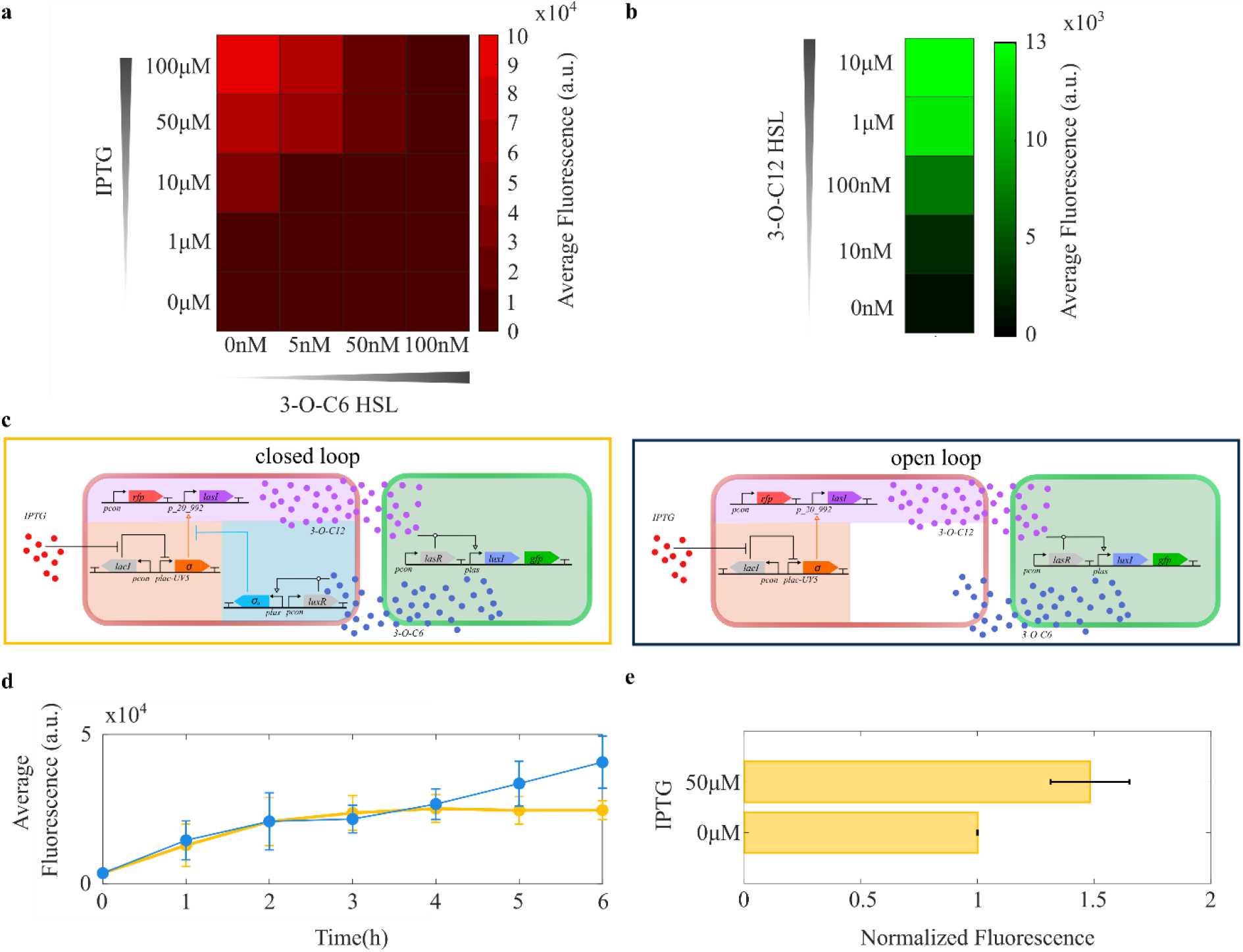
Characterization of the controllers and targets cell populations. **a:** Average fluorescence of the controller population induced using different concentrations of 3-O-C6-HSL and IPTG. The data were collected after 3h from the initial inoculation. **b:** Average fluorescence levels of the target population induced using 3-O-C12-HSL. The data were collected after 3h from the initial inoculation. **c:** Schematic representation of the closed loop (left) and the open loop (right) configurations. **d:** Average fluorescence of the targets over a 6h time course in open (blue) and closed (yellow) loop. The solid dots and vertical bars represent the average and standard deviation over n=3 biological replicates, respectively. **e:** Normalized average fluorescence of the targets mixed with the controllers in closed loop when 0 *μ*M and 50 *μ*M IPTG were present in the growth media. The data were collected 6h after the initial inoculation. The fluorescence was normalized to the average fluorescence reached when no IPTG was added in the media. All data are averaged over n=3 biological replicates.

We then assayed the ability of the two populations to communicate with each other. To test the ability of the controllers to stimulate the targets in the absence of any feedback from the latter, we implemented an open loop configuration of the consortium where the output sensing module was removed from the controller cells (see Figure 2c, right panel). This made them unable to receive feedback sensing molecules from the targets. We will refer to this population as *open loop controllers*.

We mixed the open loop controllers with the targets every hour, grew the consortium at 37 °C for 6h and analysed the targets fluorescence via flow cytometry (for further details on the experimental protocol see section *Time-course assays* of the methods). This time course, reported in Figure 2d (light blue trajectory), shows that the open loop controllers are able to communicate with the targets, i.e. that the 3-O-C12-HSL is successfully produced by the open loop controllers and sensed by the targets. Indeed, the controllers were able to induce a 15 fold increase in the average fluorescence of the targets after 6h. However, over this time period the open loop controllers were not able to stabilize the *gfp* expression levels in the targets confirming that, as expected, they were unable to receive any feedback signal sent from them. We performed similar experiments mixing the controllers with the targets. In closed loop, where controllers possess the module to sense the expression level of the controlled gene in the targets, stabilization of the targets fluorescence was observed after 3h (Figure 2d, yellow trajectory). This confirms that the targets were able to successfully produce 3-O-C6-HSL molecules which were correctly received by the controller population.

We then tested the ability of the controllers in the closed loop configuration to regulate *gfp* expression levels in the targets to different values by modulating the concentration of IPTG in the culture medium. Specifically, we grew controllers and targets in coculture at 37 °C for 6h, both with and without IPTG in the growth medium and then analysed the fluorescence level of the targets using flow cytometry. When the consortium was induced with 50 *μ*M IPTG, the relative increase in fluorescence was approximately 50% (Figure 2e). This increase was surprisingly small compared to the expected increase in the output observed during the preliminary characterisation of the populations. Comparing the 6h timepoint of the closed loop time course without IPTG shown in Figure 2d with the uninduced targets response shown in Figure 2b, we observed that the targets were expressing high levels of fluorescence when mixed with the controllers, even when no IPTG was present in the culture media (Figure S3 shows a comparison of the distribution of fluorescence of the targets in the two scenarios). This suggests that there was an unexpected high basal production of 3-O-C12-HSL, which was not revealed during the initial characterisation of the populations shown in Figure 2a. The inability to reveal the base expression level in the initial characterisation highlights the difference between the expression level needed to detect any GFP signal and the levels of enzyme needed to produce enough signalling molecules to induce the targets.

We addressed this problem by reducing the synthesis rate of *σ* in the absence of IPTG, aiming to increase the range of GFP fluorescence levels we could regulate the targets to. Specifically, we increased the repression efficiency of *plac-UV5* by LacI by adding an extra lac operator upstream of the promoter. This, as shown in [30], reduces the basal expression of the gene controlled by LacI enabling the LacI tetramer to bind two operators simultaneously, thus increasing the apparent affinity of the interaction (see *strains and constructs* section of the methods for more details on the construction of the plasmid). We will henceforth denote the modified controllers/open loop controllers “controllers/open loop controllers 2.0”. After adding the extra operator, the steady state fluorescence of the targets in closed loop and open loop was reduced to levels that were lower than the ones reached by uninduced target monocultures after 6h of growth (see Figure 3a,b). This indicated that the addition of the extra operator reduced dramatically the leakiness of the *plac-UV5* promoter. We then repeated the dynamic characterisation of the open and closed loop configurations of the consortium using the new version of both controller populations. In open loop, the open loop controllers 2.0 were able to stabilize *gfp* expression levels in the targets after 6h. In addition, by increasing the IPTG concentration in the culture media, we could achieve up to a 10-fold increase in GFP expression levels. However, the regulation was highly variable across the different biological replicates, especially at 0 *μM* and 3 *μ*M IPTG (see Figure 3a). By closing the feedback loop with the addition of the output sensing module in the controllers, we were able to increase the reliability of the regulation, reducing the standard deviation at the 6h timepoint 5-fold and 29-fold, when we used 0 *μ*M and 3 *μ*M IPTG, respectively. In addition, the settling time was comparable to the open loop configuration (approximately 3h), and we were able to induce up to a 5-fold increase when 7 *μ*M IPTG was added to the culture medium (see Figure 3b).

**Figure 3.**
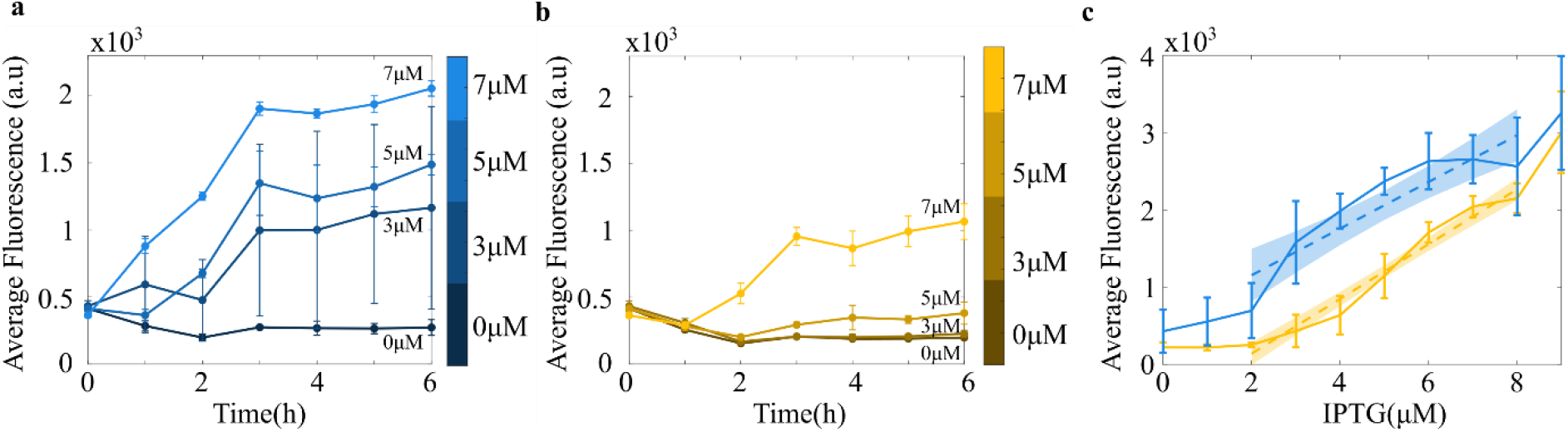
Regulation of the targets fluorescence using open and closed loop control using controllers2.0/open loop controllers 2.0. **a**. Average fluorescence of the targets over a 6h time course, using an open loop control architecture. The solid dots represent the average and the vertical bars the standard deviation of the data over n=3 biological replicates. Each trajectory is color coded and labelled with respect to the concentration of IPTG used in the experiment. **b**. Average fluorescence of the targets over a 6h time course, using a closed loop control architecture. The solid dots represent the average and the vertical bars the standard deviation of the data over n=3 biological replicates. Each trajectory is color coded and labelled with respect to the concentration of IPTG used in the experiment. **c**. Average steady state fluorescence of the targets over different IPTG concentrations. The blue (open loop) and yellow (closed loop) vertical bars are centered on the average values and their amplitude represented the standard deviation of the data over n=3 biological replicates. The dashed lines represent the linear interpolation of the data obtained with IPTG concentrations in the range [2 *μM*, 8 *μM*]. The shaded areas represent the confidence intervals of the linear model prediction. All steady states are collected after 6h from the initial inoculation.

We further investigated the tunability of the regulation in both configurations by analysing the steady state response with increasing levels of IPTG (details of the protocol used can be found in the *Titration assays* section of the Methods). Specifically, we mixed the controllers/open loop controllers 2.0 with the targets and grew the consortium for 6h at 37 °C. Figure 3c shows that both consortia were able to regulate the *gfp* gene to different expression levels. Both architectures showed saturation when IPTG ∈ [0 *μM*; 2 *μM*]. Above that concentration, increases in IPTG concentrations drove an increase in the targets’ fluorescence levels. In addition, both consortia underwent an abrupt increase in the targets’ average fluorescence and in its variance when we used IPTG at concentrations above 8*μM*. Using the remaining data (IPTG ∈ [2 *μM*, 8 *μM*]), where the response of both systems was mostly linear, we observed some differences in the regulation patterns between the two configurations. The closed loop system showed an almost linear increase in GFP fluorescence, while in open loop we noticed a sharp increase in fluorescence levels between 2μM and 3μM IPTG, followed by much flatter increase in fluorescence when IPTG was present at concentrations above 5 *μM*. We quantified the linearity of the input-output response of both configurations to IPTG by fitting the data with linear least square estimators and comparing the *R*^2^ values. Using the data where IPTG ∈ [2 *μM*, 8 *μM*], the estimator fitting the closed loop data (yellow dashed line in Figure 3c) predicts almost perfectly the variance coming from the data (*R*^2^ = 0.927). In contrast, the estimator that fits the data obtained using the open loop configuration of the consortium explained just above half of such variance (*R*^2^ = 0.624).

In our experiments, we established the effectiveness of a multicellular control architecture in regulating the expression of a desired gene within the target population. Our findings underscore the significant impact of feedback incorporation, leading to a substantial reduction in response variability and enhanced linearity in the input-output relationship of the architecture.

### Closed loop control enhances robustness to changes in the consortium composition

A crucial problem when multicellular strategies are deployed in a consortium across multiple cellular populations [31] is that different populations may grow at different rates because of differences in their metabolic loads. This may hinder the ability of the controllers to steer the dynamics of the targets as it might cause fluctuations in the production rate of the sensing (3-O-C6-HSL) and actuation (3-O-C12-HSL) molecules. We therefore analysed the robustness of both the closed loop and the open loop configurations to changes in the consortium composition (see Figure 4).

**Figure 4.**
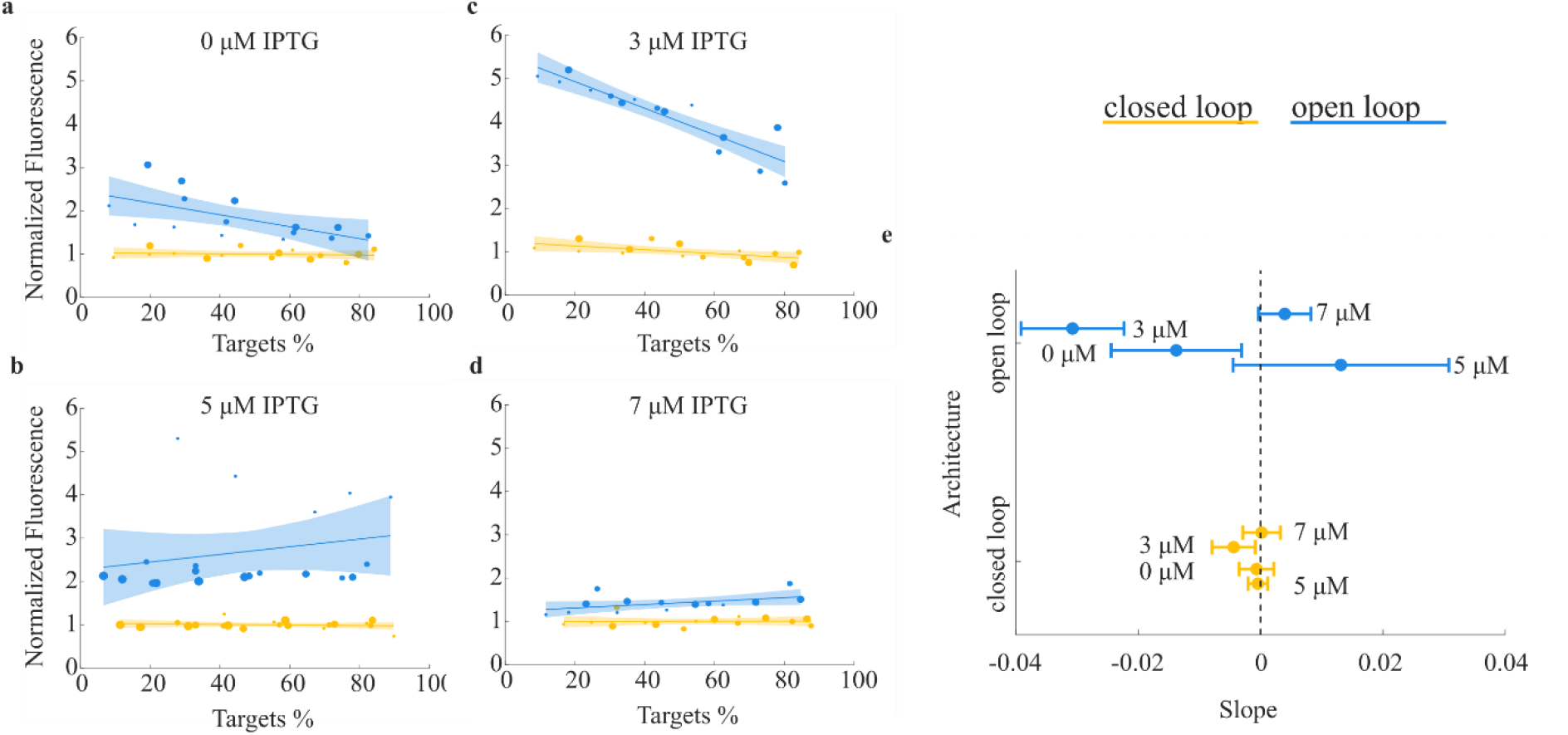
Closed loop control robustness to changes consortium composition. **a-d**. Normalized average fluorescence in open (blue) and closed (yellow) loop across different consortium compositions using controllers 2.0. /open loop controllers 2.0.. The solid dots are the data collected over n=3 biological replicates. Dots of the same size belong to the same replicate. The data were fitted with a first order polynomial. The fitting and its confidence intervals are represented with the solid lines and the shaded areas, respectively. For each replicate, the data are normalized to the average fluorescence reached in closed loop across all compositions. **e**. Comparison of the estimates of the slopes of the linear fittings of data in panels **a-d** when a closed (yellow) or open (blue) loop architecture was used. The solid dots represent the slope estimates and the horizontal bars the 90% confidence interval on the estimates.

We produced consortia of different compositions by inoculating targets and controllers/open loop controllers 2.0. in different ratios, then measuring the average fluorescence of the targets after 6h (for more detail see section *Consortium composition assay* of the Methods). Figure 4 a-d shows that varying the proportion of controllers2.0. and targets had no detectable effect on the reliability of the regulation of the closed loop configuration, where feedback between the two populations is present. Conversely, in the absence of feedback (in the open-loop configuration), we noted a declining trend in the fluorescence levels of the targets as the percentage of targets increased, especially when 0 *μ*M and 3 *μ*M IPTG concentrations were employed. This trend was not seen when 5 *μM* and 7 *μ*M IPTG were present in the culture medium. We quantified the difference between the fluorescence of the targets depending on their percentage in the consortium by fitting linear least square estimators to the data (dashed lines in Figure 4 a-d). We then compared the slopes of the lines fitting the open and closed loop datapoints at different IPTG concentrations (see Figure 4 e). Using analysis of covariance and a multiple comparison test, we found that when 0μM and 3μM IPTG were used, the slopes of the linear models fitting the closed and open loop datapoints were statistically different with *p*-values equal to 0.01527 and 1.13 10^−6^, respectively. In contrast, when 5μM and 7μM IPTG were present in the culture media, the changes were not significant. This could be attributed to the saturation of the *plas* promoter activity at higher IPTG concentrations, which reduced the sensitivity of the targets to variations in the 3-O-C12-HSL concentration (see Supplementary Figure S4 for more details and section S1 of the Supplementary Information for the details on the protocol used).

The analysis was finalized by conducting a statistical comparison of the slopes of the curves fitted to the closed-loop data points across all IPTG concentrations. Employing the same methodology, we established that there were no significant differences in slopes within the closed-loop system (detailed p-values provided in Table S3). This confirmed the consistent robustness of the multicellular feedback control strategy in maintaining stability, even in the presence of imbalances between the controllers’ and targets’ populations.

## Discussion

Our study presents for the first time the *in-vivo* biological implementation of synthetic feedback controllers within cellular populations, using a multicellular control architecture where, withing a consortium, a specific population of controller cells possesses the ability to sense and regulate targeted phenotypes within another population of cells. This strategic distribution of control functionalities across members of a microbial consortium alleviates the metabolic burden that would otherwise be imposed if all functions were concentrated within a single cell, as noted in prior research [32]. Moreover, this approach enhances the consortium’s robustness, enabling it to adapt effectively to changes in its composition.

Following the validation of individual components within our microbial community, we meticulously assessed the architecture’s performance and robustness using flow cytometry. Our results unequivocally demonstrate the distributed feedback controller’s capability to effectively regulate gene expression within target cells, maintaining the desired levels even in the face of alterations in the consortium’s composition. To substantiate the effectiveness of our control architecture, we conducted a comparative analysis between open and closed loop implementations. This examination clearly underscores the pivotal role of feedback mechanisms, revealing that their incorporation significantly enhances both the reliability and robustness of gene regulation within the system.

The development of distributed feedback controllers across microbial communities paves the way for the construction of complex interacting communities, where each population carries out a specific function [21]. Implementation of our distributed consortia within microfluidics/microscopy platforms would allow the realisation of more complex feedback control tasks in space and time [26], [33], [34].

Such microbes could be used in applications ranging from the integration of bacteria in the human gut microbiome to treat a class of diseases [35] to engineering the soil microbiome to increase plant growth and health [36], and bacteria-based production of fuels, which presents a promising source for sustainable energy [37].

Current theoretical work is aimed at improving the performance and robustness of the proposed design by expanding the consortium to include additional controller populations, adding, for example, populations able to implement proportional or derivative control actions in order to achieve a multicellular implementation of biomolecular PID controllers recently proposed in [7], [15], [38].

## Methods

### Strains and constructs

*Escherichia coli* strain TOP10 was used for all the cloning manipulations of this work. *Escherichia coli* strain MG1655 λ-, rph-1 [39] was used for all the assays in this work and transformed with the plasmids constituting either the controllers or the targets. For all the sequences of the primers and the synthesised genes used in this study see Table S1, Table S2. In addition, all the maps of the plasmids are reported In Figure S5.

The controllers contain three plasmids: pVRa20_992_BS2 (medium copy number), pVRc20_992_BS1 (medium copy number) and pVRb_LasI (low copy number). The genes constituting the essential core of the computation module (*σ*, anti-*σ* and lasI) are ssrA tagged (AANDENYALAA) to ensure fast dynamics of expression and degradation [28].

pVRb_lasI encodes the *lasI* gene, under the control of the *p_20992* promoter, and was built in [15], together with pVRb_ssrA, where the lasI gene was swapped with a superfolder *gfp*. pVRb_lasI was present in the controllers in each of the assays where they were mixed with the targets, whereas pVRb_ssra was present in the controllers for their input output characterisation.

Vector backbones for pVRa20_992_BS2 and pVRc20_992_BS1 were derived from plasmids pVRa20_992 and pVRc20_992 [27]. pVRa20_992_BS2 produces a *σ* factor under the control of a promoter that responds to IPTG.. To construct this plasmid, first the pVRa20_992_BS1 plasmid was obtained by standard restriction digestion and ligation procedure, by amplifying the *σ* gene from the pLusB plasmid [25] with primers Sigma_plus_tag_F and Sigma_plus_tag_R and cloning the PCR product in the pVRa_20_992 plasmid after NcoI/BamHI digestion. Then, the *plac* promoter was substituted with *plac-UV5* to obtain the pVRa20_992_BS2 plasmid. The construct was built by standard restriction digestion and ligation procedure, by amplifying *plac-UV5* from the pVRa_placASb_Flag plasmid [25] with primers PlacUV5_F and PlacUV5_R and cloning the PCR product in pVRa_20_992_BS1 plasmid after XbaI digestion. Finally, we modified pVRa_20_992_BS2 by introducing a second lac operator upstream of *plac_UV5*. This construct, denoted as PVRa_20_992_DS, was constructed by Gibson assembly after PCR amplification of pVRa20_992_BS2 using primers Back_extralac_for and Back_extralac_rev, and of the pVRbLacO1O1Del from [40] using the Ins_extralac_for and Ins_extralac_rev primers. pVRc20_992_BS1 produces the anti-*σ* factor under the induction of 3-O-C6-HSL. The plasmid was constructed by standard restriction digestion and ligation procedure, by amplifying anti-*σ* from pVRa_placASb_Flag [25] with primers Anti_Sigma_plus_tag_F and Anti_Sigma_plus_tag_R and cloning the PCR product in pVRc_20_992 plasmid after BamHI/PstI digestion. Finally, the controllers expressing a Red Fluorescent Protein were implemented by substituting the pVRb_lasI plasmid with the pVRb_lasI_RFP plasmid. pVRb_lasI_RFP was assembled using standard restriction digestion and ligation procedure, by amplifying a synthetically generated RFP with primers RFP_Afe_for and RFP_Nde_rev and cloning the PCR product in the pVRb_lasI plasmid after AfeI/NdeI digestion.

The targets hosted p_Las_Lux_GFP_3.0, a plasmid that embeds the 3-O-C12-HSL inducible promoter *plas*, driving the expression of a mut3 green fluorescent protein and of the luxI gene (fused with and ssrA tag). This plasmid was assembled by Gibson assembly after PCR amplification of the las_composite_device plasmid from [41] with p_Las_Lux_GFP_3_0_fwd_20 and p_Las_Lux_GFP_3_0_rev_20 primers, and of the luxI from pTeLu_GFP constructed in [25] with the luxI_fwd and luxI_rev primers.

### Chemicals

For all experiments, kanamycin (50 μg/mL), ampicillin (100 μg/mL) and chloramphenicol (25 μg/mL) were added to the growth media, depending on the bacteria selective resistance. These antibiotics were supplied by Sigma-Aldrich. pVRa_20992_BS2 and pVRa_20_992_DS have an

Ampicillin resistant gene, pVRb_lasI, pVRB_lasI_RFP and pVRb_ssrA have a gene conferring Kanamycin resistantce, and pVRc_20_992_BS1 and p_Las_Lux_GFP_3.0 have a gene guaranteeing Chroranmphenicol resistance. When multiple populations were mixed, the antibiotics to which both populations were resistant were used. If no resistance was in common, no antibiotic was used.

3-O-C6-HSL (N-(B-Ketocaproyl)-L-Homoserine Lactone from Sigma-Aldrich, cat # K3007) and 3-O-C12-HSL (N-(3-Oxododecanoyl)-L-homoserine lactone from Sigma-Aldrich, cat # O9139) were dissolved in DMSO, filter-sterilized and added to the LB growth medium at the indicated concentrations. IPTG (Isopropyl β-D-1-thiogalactopyranoside, supplied by Sigma-Aldrich, cat # I5502) was dissolved in water, filter-sterilized and added to LB medium at the indicated concentrations.

### Samples preparation and analysis using flow cytometry

Each sample (0.5 mL volume) was resuspended in 1xPBS and diluted to achieve a final OD_600_ for the culture of 0.05. Data were acquired using either BD LSRFortessa X-20 or Acea Novocyte flow cytometer. The results analysed using FlowJo.

The events recorded by a flow cytometer were first gated in the forward scatter (FSC-H, proxy of cellular size), side scatter (SSC-H, proxy of cellular complexity) plane. Here, the events were divided in three subpopulations: healthy cells where size and complexity were in physiological ranges [42], stressed cells where the bacteria had increased size and complexity due to filamentation, and cellular debris corresponding to events with high size with respect to their complexity (Figure S6 a). Subsequently, healthy cells were gated in the SSC-H, SSC-A plane to select for single cells. Cells with comparable signals in the two channels were selected as single cells, whereas events with much larger SSC-A were classified as cellular conglomerates and excluded from the analysis (Figure S6b). Finally, single cells were gated using the FITC-H (Novocyte machine)/FITC-A (Fortessa machine) (490 nm excitation, 530/30 nm emission) channel to discriminate targets and controllers based on their fluorescence levels. Specifically, cells with low FITC-H signal were classified as controllers and cells having high FITC-H fluorescence were selected to be targets (Figure S6c). Alternatively, when controllers constitutively expressed RFP, events with a high signal in the PE-CF594 (560 nm excitation, 610/20 nm emission filter, Red) channel were distinguished as controllers while cells with low signal were designated as targets.

### Induction protocol

Cells were cultured in 10 mL LB broth in 50 mL Falcon tubes. 20*μL* of overnight cultures were added to the media, which was supplemented with the inducer(s) at the specified concentration. The culture was then incubated at 37 °*C*, shaking at 250rpm for 3h. Then, the samples were prepared and analysed by flow cytometry. This protocol was used to obtain Figures 2 a,b.

### Time-course assays

35 *μL* of overnight cultures of controllers and 140 *μL* of overnight cultures of targets were separately diluted each in 35 mL LB added to a 250mL sterile flask. Each culture was supplemented with the specified concentration of IPTG. Both cultures were incubated at 37 °C, shaking at 250rpm for 6h. Each hour, 5 mL of both cultures were mixed in sterile 50 mL tubes, which were then incubated alongside the single strain cultures. After 6h, 0.5 mL of all mixed cultures were sampled and analysed via flow cytometry. Note that the hours in Figure 2c and Figure 3a,b refer to the time targets and controllers have been mixed for. Figure 2c Figure 3a,b were obtained using this protocol.

### Titration assay

For each condition, 8*μL* of overnight culture of targets and 2*μL* of overnight culture of controllers were mixed in 10 mL LB broth in 50 mL Falcon tubes. Each culture was supplemented with IPTG at the specified concentration. The cultures were then incubated at 37 °*C*, shaking at 250rpm for 6h. Then, the samples were prepared and analysed by flow cytometry. This protocol was used to obtain Figure 3 c.

### Consortium composition assay

Overnight cultures of targets and controllers were diluted in 10 mL LB using different targets:controllers ratios. Specifically, 15:1, 8:1, 4:1, 2:1, 1:1 ratios were created by adding 10*μL* of controllers and 15 0 *μL*, 8 0 *μL*, 4 0 *μL*, 2 0 *μL*, 1 0 *μL* of targets, respectively. All cultures were incubated at 37 °C shaking at 250rpm for 6h. Then, 0.5 mL of each culture was sampled and analysed by flow cytometry. This protocol was used in Figure 2d, Figure 4a-d.

## Data availability

The authors declare that the data supporting the findings of this study are available within the paper and its Supplementary Information files or from the corresponding author on reasonable request. The source data underlying all figures in the main text and supplementary information are provided as a Source Data file.

## Acknowledgements

The authors wish to acknowledge Gwen Brouwer and Abigail Smith for their support with the cloning of the constructs needed for this work. Also, the authors also wish to acknowledge the FACS facility hosted at the University of Bristol for their support with the use of the Flow Cytometers. M.d.B., L.M. wish to acknowledge support from the European Project FET-OPEN Cosy-Bio (Grant Agreement number 766840). M.d.B. also wishes to acknowledge support from the PNRR and PNC projects of the Italian Government. L.M. acknowledges funding from the Engineering and Physical Sciences Research Council (EPSRC, Fellowship EP/S01876X/1) and the Biotechnology and Biological Sciences Research Council (BrisEngBio grant, BB/W013959/1). Finally, L.M., N.J.S., C.G. and M.d.B. wish to acknowledge support form BrisSynBio, a BBSRC/EPSRC Synthetic Biology Research Centre (BB/L01386X/1).

## Author Contributions

M.d.B, L.M., N.J.S. and C.G. designed the research; D.S. designed the experiments; D.S. and B.Scarried out the experiments; D.S. with support from M.d.B., L.M., N.J.S. and C.G. analyzed the data; D.S. and M.d.B. wrote the manuscript with inputs from L.M., N.J.S. and C.G.

