## Supplementary Information for "*In-vivo* distributed multicellular control of gene expression in microbial consortia"

**Index**

S1. Sensitivity analysis to 3-O-C12-HSL in open loop……………………………………………..….. 2

SUPPLEMENTARY TABLES…………………………………………………………………….. 3

SUPPLEMENTARY FIGURES……………………………..……………………………….. 5

**S1. Sensitivity analysis to 3-O-C12-HSL in open loop**

To assess the sensitivity of the *plas* promoter to 3-O-C12-HSL when the targets were mixed with the controllers in open loop, we added exogenous 3-O-C12-HSL to the consortium (see Figure S4). The results show no statistically significant change in the average fluorescence expressed by the targets when 5 $\mu M$ and 7 $\mu$M IPTG were present. Instead, at lower IPTG concentrations, adding exogenously the actuation molecule caused a statistically significant increase in the fluorescence expressed by the targets. We also quantified the statistical relevance of the relative increase in target fluorescence levels when 1 nM 3-O-C12 was added to the media, confirming that at lower IPTG concentrations, i.e. 0 $\mu M$ and 3 $\mu M$, the increase was statistically significant (p-values 0.0008, 0.01 for 0 $\mu$M, 3 $\mu$M IPTG using a t-test). Instead, at higher IPTG concentrations, the changes were found not to be statistically relevant (p-values 0.06, 0.12 for 5 $\mu$M, 7 $\mu$M IPTG using a t-test).

### SUPPLEMENTARY TABLES

| Primer Name | Primer sequence (5′- 3′) |
|---|---|
| **Sigma_plus_tag_F** | GAGGAGAAATACCATAATCC |
| **Sigma_plus_tag_R** | GTTTTATCAGACCGCTTC |
| **placUV5_F** | TGGTTTCACATTCACCAC |
| **placUV5_R** | GCTAGATCTAGAGAAATTGTTATCCGCTCACA |
| **Anti_Sigma_plus_tag_F** | ATGTCATATGCCGCGATAGAGGAGGTAA |
| **Anti_Sigma_plus_tag_R** | CTCTCATCCGCCAAAACA |
| **Back_extralac_for** | CACCCCAGGCTTTA |
| **Back_extralac_rev** | ATTCGGGATCGAGAT |
| **Ins_extralac_for** | GAATTCCGATTCATTAATG |
| **Ins_extralac_rev** | CGGAAGCATAAAGTGT |
| **RFP_Afe_for** | TGATCGAGCGCTGATAAGTCCCTAACTTTTACAGC |
| **RFP_Nde_rev** | TGCTCACATATGGGACCAAAACGAAAAAAGGC |
| **p_Las_Lux_GFP_3_0_fwd_20** | ACGCCCTTGCAGCGTAATAATACTAGAGAAAGAG GAGAAATACTAGATGCG |
| **p_Las_Lux_GFP_3_0_rev_20** | CTCTATCGCGGAAATTGACACTCGGCGTTATGTC ATGAAG |
| **luxI_for** | TGTCAATTTCCGCGATAGAGGAG |
| **luxI_rev** | TTATTACGCTGCAAGGGCGTA |

Table S1 Name and sequence of the primers used for the cloning of the constructs used in this study.

| **Gene Name** | **Sequence (5′ – 3′)** |
|---|---|
| **RFP** | GATAAGTCCCTAACTTTTACAGCTAGCTCAGTCCAGGTATTATGC TAGCCTGAAGCTGTCACCGGATGTGCTTTCCGGTCTGATGAGTCC GTGAGGACGAAACAGCCTCTACAAATAATTTTGTTTAATACTAGA GAAAGAGGGGAAATACTAGATGGTTTCCAAGGGCGAGGAGGAT AACATGGCTATCATTAAAGAGTTCATGCGCTTCAAAGTTCACATG GAGGGTTCTGTTAACGGTCACGAGTTCGAGATCGAAGGCGAAGG CGAGGGCCGTCCGTATGAAGGCACCCAGACCGCCAAACTGAAA |

| | |
|---|---|
| | GTGACTAAAGGCGGCCCGCTGCCTTTTGCGTGGGACATCCTGAGC<br>CCGCAATTTATGTACGGTTCTAAAGCGTATGTTAAACACCCAGCG<br>GATATCCCGGACTATCTGAAGCTGTCTTTTCCGGAAGGTTTCAAG<br>TGGGAACGCGTAATGAATTTTGAAGATGGTGGTGTCGTGACCGTC<br>ACTCAGGACTCCTCCCTGCAGGATGGCGAGTTCATCTATAAAGTT<br>AAACTGCGTGGTACTAATTTTCCATCTGATGGCCCGGTGATGCAG<br>AAAAAGACGATGGGTTGGGAGGCGTCTAGCGAACGCATGTATCC<br>GGAAGATGGTGCGCTGAAAGGCGAAATTAAACAGCGCCTGAAA<br>CTGAAAGATGGCGGCCATTATGACGCTGAAGTGAAAACCACGTA<br>CAAAGCCAAGAAACCTGTGCAGCTGCCTGGCGCGTACAATGTGA<br>ATATTAAACTGGACATCACCTCTCATAATGAAGATTATACGATCG<br>TAGAGCAATATGAGCGCGCGGAGGGTCGTCATTCTACCGGTGGC<br>ATGGATGAGCTGTACAAATAACTCGGTACCAAATTCCAGAAAAG<br>AGGCCTCCCGAAAGGGGGGCCTTTTTTCGT TTTGGTCC |

Table S2 Names and sequences of the gene commercially synthesised for this study.

| IPTG CONCENTRATION | 0 $\mu$M | 3 $\mu$M | 5 $\mu$M | 7 $\mu$M |
|---|---|---|---|---|
| 0 $\mu$M | - | 0.219 | 1.000 | 0.973 |
| 3 $\mu$M | 0.219 | - | 0.179 | 0.098 |
| 5 $\mu$M | 1.000 | 0.179 | - | 0.965 |
| 7 $\mu$M | 0.973 | 0.098 | 0.965 | - |

Table S3 p-values of a multiple comparison analysis run on the slope of the linear least square estimators fitted on closed loop data spanning different consortium compositions Details on how the data were obtained can be found in section Consortium composition assays of the Methods.

SUPPLEMENTARY FIGURES

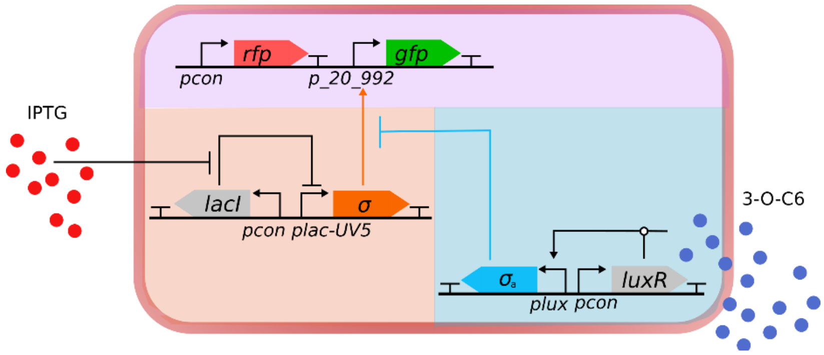


Figure S1 Schematic representation of the controllers used to characterise the input-output response. In this population the plasmid expressing lasI under the p_20_992 promoter was modified substituting lasI with a green fluorescent protein (GFP).

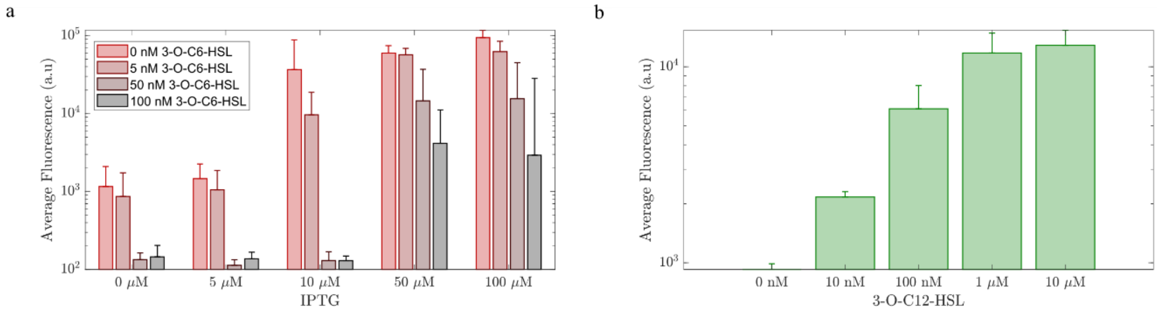


Figure S2 Steady state response of controllers and targets In single strain cultures. **a**. Average fluorescence of the controller population induced using different concentrations of 3-O-C6-HSL and IPTG. The data were collected 3h after the initial inoculation. The bar and vertical lines depict the average and standard deviation of the fluorescence expressed by the controllers over n=3 biological replicates. **b**. Average fluorescence of the targets induced using different concentrations of 3-O-C12-HSL. The data were collected 3h after the initial inoculation. The bar and vertical lines depict the average and standard deviation of the fluorescence expressed by the fluorescent controllers over n=3 biological replicates.

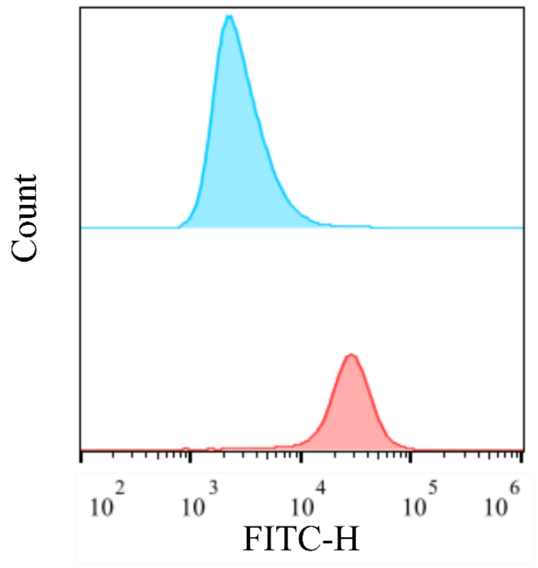


Figure S3 Fluorescence profile of the targets in monostrain culture (blue) and in co-culture with the controllers (red) 6h after the initial inoculation. In both conditions no IPTG or 3-O-C12 were added to the growth medium.

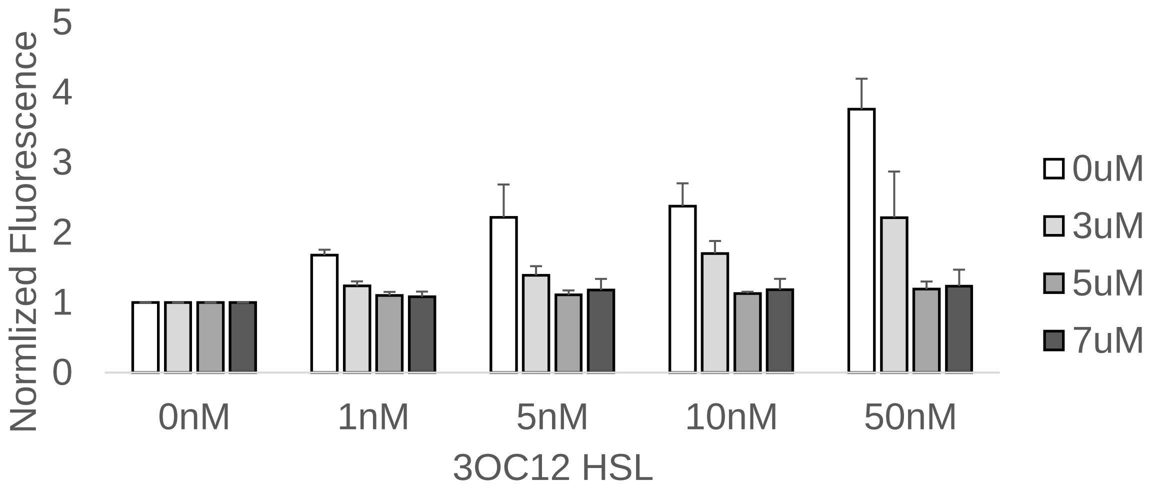


Figure S4 Normalized fluorescence levels of the target population when increasing levels of 3-O-C12 HSL were added to the culture media. The targets were mixed with the open loop controllers 2.0. and induced using different IPTG concentrations. The data were collected after a 6h incubation at 37 $^{o}$C, shaking at 250rpm. Each IPTG condition was normalized to the case where no 3-O-C12 HSL was present in the medium.

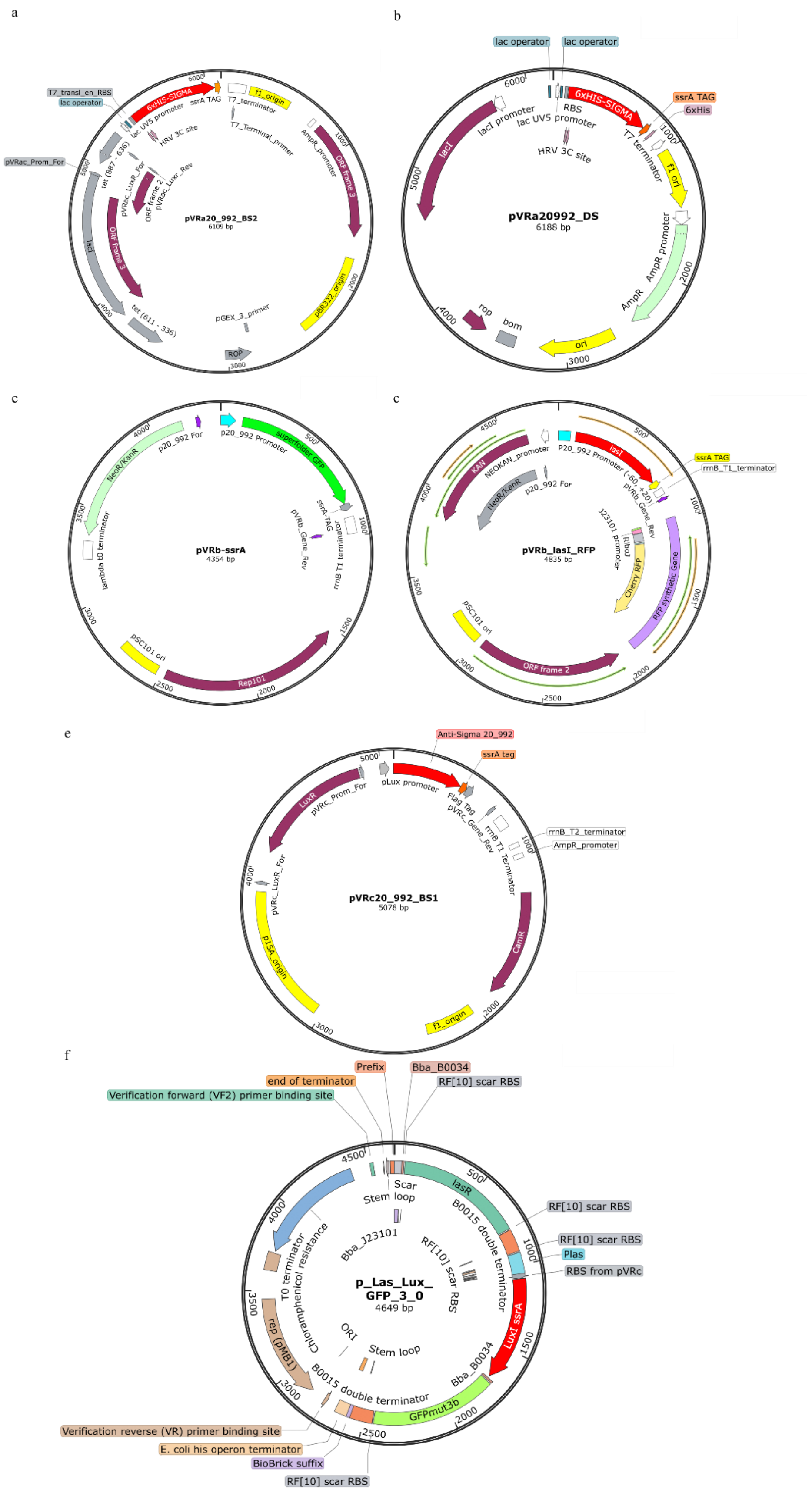


Figure S5 Plasmid maps of the construct used in this study. **a** map of the pVRa_20_992_BS2 ($\sigma$ producing plasmid of the controllers) **b** map of the pVRa_20_992_DS ($\sigma$ producing plasmid of the controllers 2.0) **c** map of the pVRb_ssrA (plasmid producing GFP under the p_20_992 promoter) **d** map of the pVRb_lasI_RFP (plasmid producing lasI under the p_20_992 promoter) **e** map of the pVRc_20_992_BS1 (plasmid producing anti-$\sigma$) **f** map of the plas_lux_GFP_3.0 (plasmid embedded in the targets)

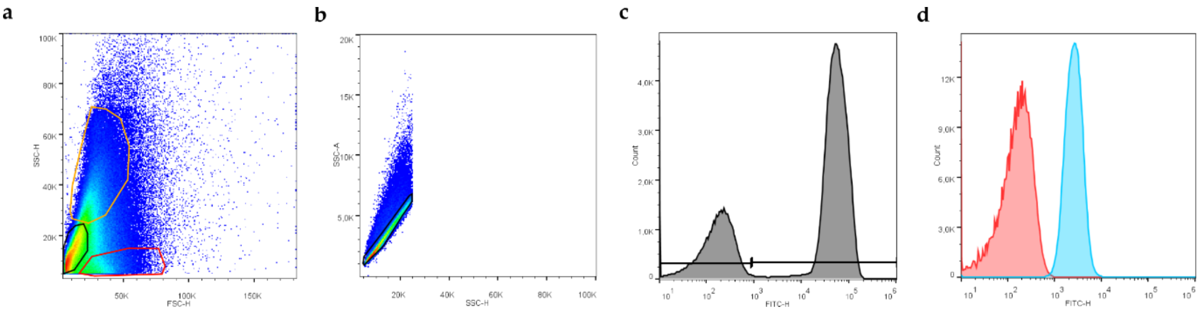


Figure S6 **a.** Density plot of events in the FSC-H,SSC-H plane. Red areas represent zones of the plane with more events. The Polygons drawn represent events classified as healthy cells (black), stressed cells (orange) and cellular debris (red). **b.** Density plot healthy cells in the SSC-H,SSC-A plane. Red areas represent zones of the plane with more events. The black polygon drawn is the gate used to select single cells against aggregates. **c.** Histogram of single cells in the FITC-H (Green Fluorescence) channel. The two horizontal gates highlights controllers and targets in the consortium. **d.** Comparison of the FITC-h fluorescent profiles of controllers (red) and uninduced targets (blue).
